# Efficient Estimation of Nucleotide Diversity and Divergence using Depth Information

**DOI:** 10.1101/2025.05.15.654353

**Authors:** Cade Mirchandani, Erik Enbody, Timothy B. Sackton, Russ Corbett-Detig

## Abstract

The increasing scale of population genomic datasets presents computational challenges in estimating summary statistics such as nucleotide diversity (π) and divergence (*d*_xy_). Unbiased estimates of diversity require knowledge of missing data and existing tools require all-sites VCFs. However, generating these files is computationally expensive for large datasets. Here, we introduce **Callable Loci And More (clam)**, a tool that leverages callable loci—determined from depth information—to estimate population genetic statistics using a variant-only VCF. This approach offers improvements in storage footprint and computational performance compared to contemporary methods. We benchmark *clam* using a large muskox dataset and demonstrate that it produces unbiased estimates of π while reducing runtime and storage requirements, compared to an existing approach. *clam* provides an efficient and scalable alternative for population genomic analyses, facilitating the study of increasingly large and diverse datasets. *clam* is available as a standalone program and integrated into snpArcher for efficient reproducible population genomic analysis.

## Introduction

In recent years, rapid advancements in sequencing technology have dramatically lowered the cost of genome sequencing, enabling the generation of numerous, large population resequencing datasets (Corbett-Detig et al. 2015). However, the volume of data now available (All of Us Research Program Genomics Investigators 2024) presents new computational challenges. To analyze large datasets effectively, there is a pressing need for tools that can efficiently handle the scale and complexity of high-throughput genomic data while maintaining analytical accuracy (Hendricks et al. 2018).

Nucleotide diversity (π), divergence (*d*_xy_), heterozygosity, and *F*_ST_ are fundamental population genetic summary statistics that summarize key aspects of population history and structure. π measures genetic variation within populations (Nei and Li 1979), *d*_xy_ quantifies genetic distance between populations (Nei and Li 1979), heterozygosity reflects genetic diversity at the individual level, and *F*_ST_ measures population differentiation (Hudson et al. 1992; Bhatia et al. 2013). Accurate estimation of these statistics is essential for inferring demographic history, detecting selection, and understanding speciation processes (Hendricks et al. 2018; Luikart et al. 2018; Hohenlohe et al. 2021).

These fundamental population-genetic parameters are commonly estimated using polymorphism data stored in Variant Call Format (VCF) files (Danecek et al. 2011), which typically only records genomic positions where variants were identified in the population. While this type of VCF (from here on referred to as a “variants-only VCF”) is widely accepted as the standard format for most analyses of genetic polymorphism, it poses challenges for estimating statistics that rely on pairwise nucleotide comparisons, such as π and *d*_xy_. When using a variant-only VCF to estimate these statistics, sites not included in the file are treated as homozygous reference, which can bias estimates, as the total number of sites used in the calculations is overestimated, artificially deflating the values of these statistics (Korunes and Samuk 2021).

To address this bias, an “all-sites” VCF that includes both variant and invariant positions can be used as input to calculate these statistics (Korunes and Samuk 2021). While this solution produces more accurate estimates of diversity, the creation, storage, and analysis of all-sites VCFs poses computational challenges as genomic datasets grow larger, making them impractical to produce, analyze and store. Additionally, filtering approaches for all-sites VCFs often differ from those applied to variant-only VCFs, introducing potentially unaccounted for confounding effects to population genomic analyses. One alternative approach to using an all-sites VCF, implemented in ANGSD (Korneliussen et al. 2014), uses genotype likelihoods across all sites to derive allele frequencies which are then used to estimate population genetic statistics. However, as (Korunes and Samuk 2021) points out, ANGSD lacks validated protocols to produce estimates of π or *d*_xy_, and as such there may be inconsistency in estimates across studies using this approach.

An alternative approach is to leverage a variant-only VCF and sequencing depth to determine whether each site in the genome is sufficiently covered to be considered “callable”. This information can be used to fill in the gaps of a variant-only VCF by identifying regions where sequencing coverage was sufficient to call genotypes, but are not present in the VCF, allowing these positions to be counted as invariant sites in diversity calculations. However, there are no maintained tools that both efficiently identify callable loci and use callable loci information to estimate population genetic statistics. Here, we present Callable Loci And More (*clam*), a command line tool that estimates population genetic statistics using a VCF and callable loci information. We show that *clam* produces minimally biased estimates of π, while requiring less disk space and less compute time. *clam* will enable population genetic insights from increasingly vast datasets.

## Methods

### Implementation

*clam* is a command line tool written in the Rust programming language that provides two subcommands: *loci* and *stat*. The *loci* subcommand generates callable loci interval files by analyzing per-sample sequencing depth, accepting input in the form of either per-sample depth information stored in D4 files (Hou et al. 2021) or GVCFs. D4 files are highly efficiently compressed estimates of sample depth for every site and are easily produced (e.g. using mosdepth) from alignment (i.e. bam) files. The *stat* subcommand accepts a VCF and the callable loci file as inputs to simultaneously count pairwise differences between samples at polymorphic sites in the VCF and lookup the total number of sites where genotypes could be reliably called in the callable loci file. *Stat* calculates π and *d*_xy_, *F*_ST_ (if applicable population labels are provided), and per-sample heterozygosity in genomic windows. Source code and documentation for *clam* is available here: https://github.com/cademirch/clam and can be installed either by compiling from source or via bioconda (Grüning et al. 2018).

### Empirical Dataset

To benchmark the computational requirements, we reanalyzed a publicly available population genomic dataset of muskox (*Ovibos moschatus*; N=189) (Pečnerová et al. 2024). We selected this dataset as the muskox genome is relatively large (2.6 Gb) and low heterozygosity (0.6 x10^−4^ - 2.4×10^-4^), both features that will produce very large all-sites VCFs. We first used the Snakemake workflow (Mölder et al. 2021) snpArcher (Mirchandani et al. 2024) to generate per-sample GVCFs. Briefly, we obtained sequencing reads from the SRA and ENA (Leinonen et al. 2011), then we trimmed the adapter sequence using *fastp* (Chen *et al*. 2018). We aligned the trimmed reads to the muskox reference genome using BWA-MEM (Li 2013), marked duplicate alignments using Sentieon *Dedup*, and then called individual variants (GVCF) using Sentieon *Haplotyper* (Kendig et al. 2019).

Next, using a modified version of snpArcher we performed joint genotyping per-contig (excluding contigs smaller than 100kb) using Sentieon *Genotyper* with the option “--emit_mode=VARIANT” to produce the variant only VCFs and “--emit_mode=ALL” to produce all-sites VCFs. We then split the all-sites VCFs into variant and invariant sites using vcftools (Danecek et al. 2011). We applied identical filtering criteria to both the variant sites extracted from the all-sites VCF and the variants-only VCF using *bcftools* (Danecek *et al*. 2021), removing sites that failed GATK hard filtering thresholds (McKenna et al. 2010; Van der Auwera et al. 2013), sites with minor allele frequency <0.01, and sites with >75% missing data. For invariant sites, we used bcftools to set genotypes to missing (“./.”) at positions with read depth less than 10, then concatenated the filtered variant sites and filtered invariant sites to produce the final all-sites VCF.

To generate the callable loci file, we first computed per-base depth profiles for each sample from the duplicate-marked alignments using *mosdepth* (*Pedersen and Quinlan 2018*) to produce D4 files, then used *clam loci* to count how many samples were “callable” (read depth >10) at each position in the genome. We then estimated π in 100kb windows using *clam stat* with the variants-only VCF and callable sites, and *clam stat* with the all-sites VCF. Additionally, we used *pixy* (Korunes and Samuk 2021) to estimate π in 100kb windows from the all-sites VCF.

## Results & Discussion

### Runtime comparison of all-sites and variants-only workflows

The all-sites workflow to estimate diversity required substantially more computational time than the variants only and callable loci approach, even when accounting for the additional steps of running mosdepth and *clam* loci in the latter. To approximate the total computational requirements to run each workflow, we summed the runtime of all jobs within each workflow step from joint genotyping onwards, as the preceding steps (fastq processing, alignment, and per-sample variant calling) were shared in both approaches. This summation provides a measure of computational demand of each workflow, although realized walltime will vary based on computational infrastructure. The relative difference between workflows should remain relatively consistent regardless of specific computing environment, making this comparison a reliable metric of the computational cost of each approach.

In total, the all-sites workflow required 219.6 hours of processing time, 9.25x greater than the 23.7 hours needed for the variants-only and callable loci approach (Table 1). The filtering invariant sites step in the all-sites workflow was the largest contributor to the compute time difference, requiring 152.16 hours, or 77.7% of the additional runtime required by the all-sites workflow. Joint genotyping took 8.05x longer in the all-sites workflow (45.78 hours vs 5.68 hours), accounting for 20.5% of the difference, and filtering variant sites required 3.67x more time, or 7.6% of the total difference. Additionally, the two workflows differed in their storage requirements. The all-sites approach generated 229.96 GB of VCF files, while the variants-only approach produced only 48.67 GB of VCFs plus an additional 12.26 GB callable loci file, for a total of 60.93 GB (Table 2).

**Table 1.**
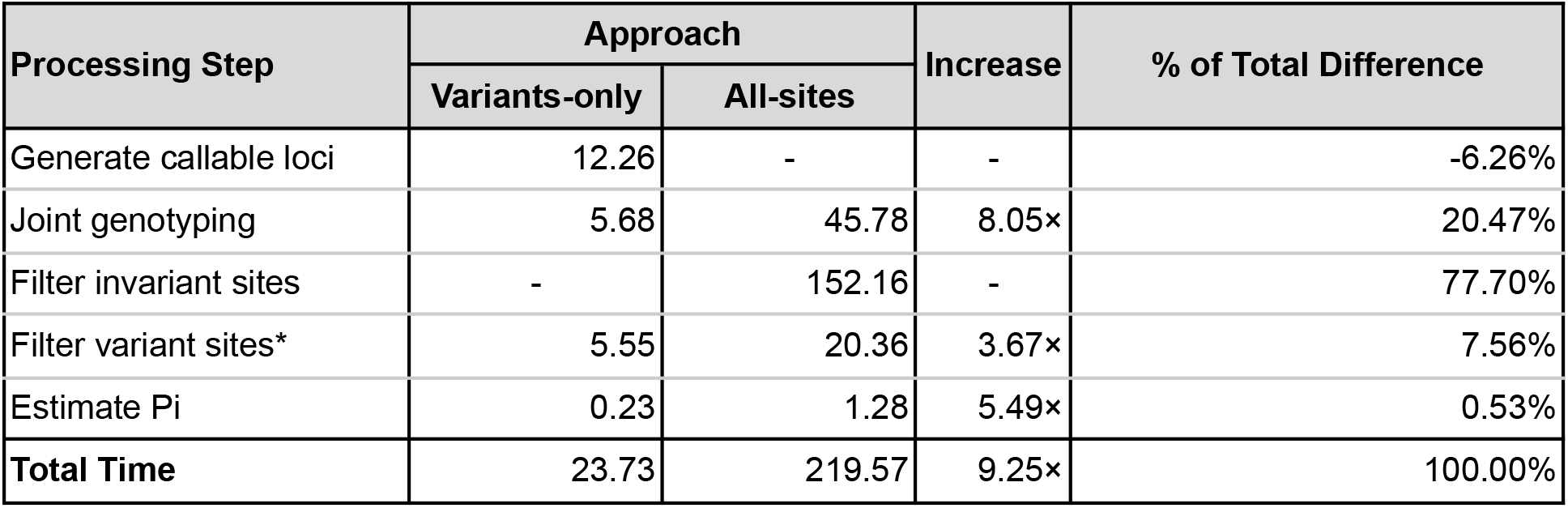
Computational runtime requirements of all-sites and variants-only workflows. Each value represents the sum of wall clock time (in hours) for all jobs within the workflow step. The “filtering variant sites” in the all-sites workflow includes time to separate variant and invariant sites.

**Table 2.** Disk storage requirements of all-sites and variants-only workflows in gigabytes.

| File Type | Approach |  | Increase |
| --- | --- | --- | --- |
|  | Variants-only | All-sites |  |
| VCF Files | 48.67 | 229.96 | 4.72× |
| Callable loci file | 12.26 | - | - |
| **Total** | 60.93 | 229.96 | 3.77× |

The substantial differences in computational demands between approaches largely stem from the increased I/O (input/output) operations required to create and analyze the all-sites VCFs, which contain many more sites than variants-only VCFs. The variants-only and callable loci approach circumvents this by using a compact representation of invariant sites in the form of callable loci intervals rather than per-base invariant VCF records. This approach reduces the volume of data that must be processed and stored, greatly reducing both computational processing time and storage requirements. The magnitude of these improvements will likely scale with genome size and number of samples, making this approach ideal for large population genomic datasets.

### *clam* accurately estimates diversity without all-sites VCFs

To validate the accuracy of *clam*, we first compared its diversity estimates to those from *pixy*, an established tool for calculating population genetic statistics. We estimated π in 100kb windows across the muskox genome using three approaches: 1) *clam stat* using the variants-only VCF and callable loci, 2) *clam stat* using the all-sites VCF, and 3) *pixy* using the all-sites VCF. *clam stat* with the all-sites VCF produces identical estimates to *pixy* (Fig 1A), verifying the underlying accuracy of the *clam* algorithm.

**Figure 1.**
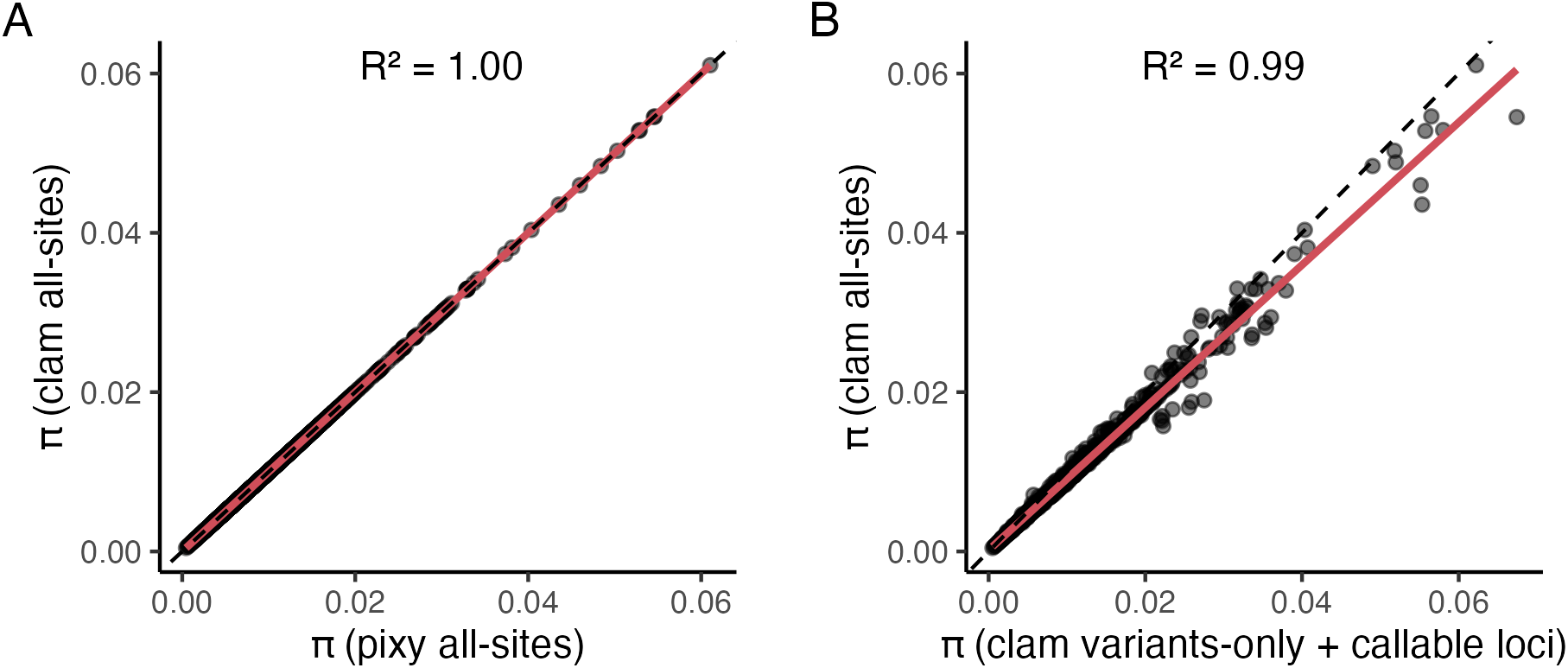
π estimated by *clam* and *pixy* using either variants-only VCF and callable loci, or all-sites VCF. A) π estimates in 100kb windows estimated by *pixy* using all-sites VCF and *clam* using the all-sites VCF. Red line shows the trend, the dashed line represents y=x. B) π estimates in 100kb windows estimated by *clam* using the callable loci approach and *clam* using the all-sites approach. Red line shows trend, dashed line represents y=x.

To demonstrate that the variants-only and callable loci approach works effectively, we compared *clam stat* results using the variants-only VCF against those using the all-sites VCF. These comparisons show that the callable loci approach produces highly similar estimates of π (R^2=0.99) to the all-sites approach (Fig 1B). We do not expect perfect 1:1 correspondence because the all-sites VCF uses VCF depth to determine missingness while the callable loci uses BAM depth, and these are not exactly equivalent due to the *de novo* assembly and local realignment performed by the variant caller to produce the VCF. Despite these technical differences in depth filtering, the high correlation between approaches demonstrates callable loci can effectively replace the need for complete all-sites VCFs when estimating diversity.

Furthermore, this highlights the effect variant filtering has on estimating population genetic statistics such as π. We also note that *clam* in itself will not completely mitigate the effect of reference bias wherein non-reference alleles are challenging to align, but methods such as *vg (Garrison et al. 2018)* in combination with this approach could help resolve this challenge.

## Conclusion

*clam* provides a computationally efficient and scalable approach for estimating population genetic statistics from large genomic datasets. By leveraging callable loci from depth information, *clam* eliminates the need for computationally expensive all-sites VCFs while maintaining unbiased estimates of nucleotide diversity (π) and divergence (*d*_xy_). Our benchmarking analysis using muskox genomic data demonstrates that *clam* significantly reduces runtime and storage requirements compared to traditional workflows. We also integrate *clam* into snpArcher, a end-end variant calling workflow written in Snakemake. This integration reduces the overhead for integrating *clam* into data analyses workflows, as variant data produced from snpArcher can easily be used for calculating population genetic statistics as described here. These advantages make *clam* particularly well-suited for studies involving massive sequencing datasets, notably those with larger genomes and low genetic diversity, where computational efficiency is critical. As genomic datasets continue to grow, *clam* offers an essential tool for researchers seeking accurate and efficient population genetic analyses.

## Data Availability

Source code and documentation for *clam* is available at https://github.com/cademirch/clam. Raw sequence reads for the muskox dataset are available in the European Nucleotide Archive under study accession ID: PRJEB64293, and reference genome available under the NCBI GenBank assembly accession ID: GCA_041156055.1.

## Acknowledgements

We are grateful for the computational resources provided by the UCSC Genomics Institute. TBS was funded by the National Science Foundation (NSF) Division of Biological Infrastructure award DEB-1754397.

## Notes

### Competing Interest Statement

The authors have declared no competing interest.

